# Xingnaojing injection can regulate dysbacteriosis and increase the concentration of short chain fatty acids in the feces after stroke in mice

**DOI:** 10.1101/2020.11.05.370528

**Authors:** Jingfeng Lin, Ganlu Liu, Zhenyun Han, Qiang Gao, Zhenyi Wang, Ze Chang, Ying Gao, Dayong Ma

**Affiliations:** Beijing University of Chinese Medicine, Beijing, 100029, China; Shenzhen Hospital of Beijing University of Chinese Medicine (Longgang), Shenzhen, 518100, China; Neurology Department of Dongzhimen Hospital, Beijing University of Chinese Medicine

**Keywords:** Xingnaojing, gut microbiota, short chain fatty acids, stroke

## Abstract

**Background:** Xingnaojing injection (XNJ) is extracted from the Chinese ancient prescription “An-Gong-Niu-Huang Pill”, is widely used for stroke in China. We mainly observe the effect of XNJ (Xingnaojing) injection on the gut microbiota in stroke model mice.

**Methods:** Forty-two 7-to 8-week-old male C57 mice weighing 22-24 g were chosen for the experiment. There were 6 mice in each group; the 7 groups were the normal group (NG), the MCAO group (CG), the MCAO+XNJ group (EG), the sham surgery group (SG), the sham germ-free normal group (SGFNG), the sham germ-free+MCAO group (SGFCG), and the sham germ-free+MCAO+XNJ group (SGFEG). Two days before modeling, we abdominally administered Xingnaojing (6 mg/kg) the SGFEG and EG groups. The processing time of sustained XNJ was 5 days. Three days after modeling, 1 ~ 2 mouse feces were collected, and after a MiSeq PE library was constructed, an Illumina MiSeq PE 300 platform was used for high-throughput sequencing. After cleaning the sequencing data, the microbiome and microbiomeseq packages were used for analysis using R software (version 3.6.2).

**Results:** Alpha diversity analysis revealed that the diversity was not different between the CG and EG. The Simpson index was different between the SGFCG and SGFEG. XNJ increased the levels of *Sutterellaceae* and decreased the level of *Deferribacteres* and *Morganella*. LEfSe analysis showed that SGFCG mice were also enriched with *Morganella*. XNJ increased the concentrations of the SCFAs PA (propionate), VA (valerate), IBA (isobutyrate), and IVA (isovalerate) in the feces of the SGFEG group. BA (butyrate) had greater positive correlation with gut bacteria than other acids in the SGFCG, and XNJ changed this trend. KEGG analysis showed that peptidoglycan biosynthesis was most different between the CG and EG.

**Conclusion:** Ischemic stroke (IS) causes dysbiosis of some specific bacteria in the gut microbiota in MCAO mice. Xingnaojing ameliorated this condition by increasing the levels of *Sutterellaceae* and decreasing the level of *Deferribacteres* and *Morganella*. These results are in accordance with other research on Chinese medicines for IS that affect the gut microbiota. Enrichment analysis of SCFAs revealed that XNJ improved the levels of SCFAs through an energy metabolism-related pathway.

## Background

Xingnaojing injection (XNJ) is extracted from the Chinese ancient prescription “An-Gong-Niu-Huang Pill”, has been approved by the China Food and Drug Administration (CFDA) and is widely used for the treatment of stroke^[1]^. XNJ is composed of four Chinese herbs. The components are shown in Table1^[2]^.

**Table 1.** Components of Xingnaojing injection

| Traditional Chinese<br>medicine name | Moschus | *Radix curcumae* | Borneol | *Fructus*<br>*gardeniae* |
| --- | --- | --- | --- | --- |
| Binomial | Not plant | *Curcuma aromatica* | Not plant | *Gardenia* |
| nomenclature names |  | *Salisb* |  | *jasmnoides J.Ellis* |
| Part | Not applicable | root | Not applicable | fruit |
| Main ingredients | Musclide B | Acetic acid | (+)-Borneol | Geniposide |
|  | Hyperin | Spathulenol |  | Epijasminoside |
|  | Gibberellin A8 | Oxalic acid |  | A |
|  | (+)-Jurinelloide | 4-Hydroxybenzoic |  | Gardaloside |
|  | Gibberellin A1 | acid |  | Acetylursolic |
|  | Gibberellin A17 | Serotonin |  | acid |
|  | Gibberellin A4 | Thymol |  |  |
|  | trans,trans-Farnesol | L-Asparagine |  |  |
|  |  | Zederone |  |  |
|  |  | Curcumenolactone C |  |  |
|  |  | Zedoarolide A |  |  |
|  |  | Zedoarolide B |  |  |
|  |  | Sucrose |  |  |
|  |  | Zerumin B |  |  |
|  |  | Curcumin |  |  |
|  |  | Curcumanggoside |  |  |
|  |  | Tryptamine |  |  |
|  |  | alpha-Pinene |  |  |
|  |  | 4-Hydroxybenzaldehy |  |  |
|  |  | de |  |  |
|  |  | 13-Hydroxygermacro |  |  |
|  |  | ne |  |  |
|  |  | Curcumlactone |  |  |
|  |  | Zedoarondiol |  |  |
| Compatibility | 7.5 | 30 | 1 | 30 |
| proportion |  |  |  |  |

\*Main ingredients were detected by Hybrid Quadrupole-TOF LC/MS/MS Mass Spectrometer, Component ID: TripleTOF 5600+, Manufacturer: AB Sciex Instruments, Shimadzu LC30

It has the functions of enlightening the brain, calming pain, clearing heat and detoxification, tranquilizing the mind and cooling the blood in Chinese medicine^[3, 4]^. XNJ is widely used for stroke in China^[5]^.

In clinical research, a meta-analysis including 53 randomized controlled trials with 4915 participants showed that XNJ might be a beneficial therapeutic agent for the treatment of cerebral infarction^[6]^. Xinxing Lai^[5]^ designed a multicenter, prospective, randomized, controlled, open-label trial with blinded endpoints, which provided critical evidence for the efficacy of XNJ administration for the treatment of acute ischemic stroke (AIS) as a complementary approach initiated after reperfusion therapy or when the AIS patients are not eligible for thrombolytic treatment. Based on previous clinical research, we hypothesized that XNJ could have certain effects against ischemic stroke (IS).

Further studies on the mechanism by which XNJ alleviates ischemic stroke have demonstrated that XNJ may have multiple neuroprotective mechanisms to improve neurobehavioral disturbances and reduce the infarct size in vitro or animal works. The mechanisms include inhibition of glutamate-induced apoptosis^[7]^, antiautophagic effects via the p53-DRAM signaling pathway ^[8]^, regulations of the phosphorylation levels of proteins associated with the PI3K/Akt/eNOS signaling pathway^[2]^ and regulation of the SIRT1-mediated inflammatory response inhibition pathway^[1]^. Rong Ma^[9]^ identified 23 eligible animal studies with totally of 578 animals (290 in the trial group; 288 in the control group) and made a meta-analysis, found that XNJ could significantly reduce the neurological deficit score, alleviate the cerebral infarction area, brain edema, and the neuronal cell apoptosis, and the mechanism of efficiency in cerebral ischemia maybe was regulating oxidative stress and inflammatory reaction. Xinhua Xia^[10]^ used the model of focal cerebral ischemia reperfusion performed in rats by MCAO method to find that the Moschus and Borneol(two main components of XNJ) could significantly ameliorate neurobehavioral disturbances, shrink relative infarct size, and rescue neural dysfunction effectively. Gang Wei^[8]^ proposed that the inhibition of autophagy via p53-DRAM signaling pathway is an important mechanism of protection after experimental stroke by XNJ via vitro experiment. Yueming Zhang^[1]^,^[2]^ found that XNJ could improve rat cerebral ischemic injury and OGD induced HBMECs apoptosis, and that might be relevant to the activation of PI3K/Akt/eNOS signaling pathway and the SIRT1-mediated inflammatory response inhibition pathway via vivo and vitro researches.

In recent years, studies have found that many diseases are associated with gut microbiota imbalance^[11]^ and that altering the gut microbiota has therapeutic potential. In addition, the gut-brain axis (GBA) has been defined, comprising a two-way communication network between the brain and the gastrointestinal tract, which links the central cognitive and emotive centers of the brain with peripheral intestinal processes^[12]^. This interaction between the gut microbiota and brain is bidirectional, involving neural, endocrine, immune, and humoral links^[13]^. Based on the above findings, we hypothesized that XNJ can regulate the gut microbiota and intestinal bacteria metabolites, which may be one of its brain protection mechanisms.

Previous studies have suggested that the intestinal microbiota may regulate trimethylamine N-oxide (TMAO) via a metabolic pathway, which could alter the risk of stroke^[14–16]^. Xiuli Zeng compared different stroke risk levels and found that opportunistic pathogens and lactate-producing bacteria were enriched in the high-risk group compared to the low-risk group, while butyrate-producing bacteria were depleted. Butyrate concentrations were also lower in the fecal samples obtained from the high-risk group than those from the low-risk group^[17]^. Some studies have also determined that stroke leads to gut microbiota dysbiosis and that the gut microbiota influences stroke outcomes^[18, 19]^. Based on changes between poststroke models and controls, Li Na ^[20]^ and Kazuo Yamashiro^[18]^ concluded that gut dysbiosis is associated with metabolism and systemic inflammation and that the gut microbiota can also influence stroke outcomes through the immune pathway^[18, 21, 22]^.

However, there have been no studies on the ability of XNJ injection to regulate the gut microbiota. To observe the effect of XNJ injection on the intestinal flora and metabolites of the gut microbiota after stroke, we designed this experiment to explain the mechanism underlying the efficacy of XNJ injection in the treatment of ischemic stroke from the perspective of intestinal microbiota. The experiment was approved by Animal Ethics Committee of Beijing University of Chinese Medicine.

## Methods

### 2.1. Animals

Forty-two 7-to 8-week-old male C57 mice were bought form company (Geneline Bioscience, Shanghai) weighing 22-24 g were chosen for the experiment. They were given free access to food and water and housed at a temperature of 23±2⎕ and a relative humidity of 40% ~ 70%. They were kept on a 12-h diurnal light cycle. Water bottles and other utensils used to feed the mice were all soaked in diluted 84 disinfectant for 30 min, rinsed with tap water, and allowed to dry naturally. Drinking water was autoclaved and brought to room temperature before it was provided to the mice. Other procedures were performed in strict adherence to the mouse room specifications.

### 2.2. Experimental design

The experiment began after 3 days of adaptive feeding. Using a computer-generated random number table, 6 mice were assigned to each group; the 7 groups were the control group (CG, MCAO group), the experiment group (EG, Xingnaojing group), the normal group (NG), the sham surgery group (SG), the sham germ-free group (SGFCG), the sham germ-free experiment group (SGFEG) and the sham germ-free normal group (SGFNG). The model was induced in the sham germ-free for 14 days. All mice were given free access to food and water. In addition, SGFCG, SGFEG and SGFNG mice were oral administered ampicillin (200 mg/kg, 1 g/L, Tenglong Medicine, Jiangxi), metronidazole (200 mg/kg, 1 g/L, Shijiazhuang No. 4 Pharmaceutical), neomycin sulfate (200 mg/kg, 1 g/L, Meidiya Biotech Company, Shanxi) and vancomycin (200 mg/kg, 0.5 g/L, LUMMY, Chongqing) once every two days over a period of 14 days. On the last two days of the 14-day period and 2 days before modeling, we injected Xingnaojing (6 mg/kg, Jemincare Medicine, Jiangxi, Chinese approval number: Z32020562) into the SGFEG and EG mice abdominally. The groups and their interventions are outlined in Table 1.

**Table 1.** The 7 groups and their interventions

| Group | Abbreviation | Intervention | Processing Time | Surgical approach |
| --- | --- | --- | --- | --- |
| Normal group | NG | Equal volume of normal saline injection(6 ml/kg) once a day abdominally | 5 days | No surgery |
| Sham surgery group | SG | Equal volume of normal saline injection(6 ml/kg) once a day abdominally | 5 days | Isolation of the artery with no occlusion |
| MCAO group (Control group) | CG | Equal volume of normal saline injection(6 ml/kg) once a day abdominally | 5 days | Middle cerebral artery occlusion (MCAO) |
| Xingnaojing group (Experiment group) | EG | Xingnaojing injection (6 ml/kg) once a day abdominally | 5 days | Middle cerebral artery occlusion (MCAO) |
| Sham germ-free experiment group | SGFEG | Xingnaojing injection (6 ml/kg) once a day abdominally | 5 days | Middle cerebral artery occlusion (MCAO) |
| Sham germ-free group | SGFCG | Equal volume of normal saline injection(6 ml/kg) once a day abdominally | 5 days | Middle cerebral artery occlusion (MCAO) |
| Sham germ-free normal group | SGFNG | Equal volume of normal saline injection(6 ml/kg) once a day abdominally | 5 days | No surgery |

The animals were injected abdominally for 5 days, from 2 days before modeling to 3 days after modeling.

### 2.3 Experimental stroke model and sample collection

Mice were anesthetized with 2% pentobarbital (45 mg/kg). A nylon suture was inserted into the right external carotid artery and advanced until it obstructed the middle cerebral artery, and the common carotid artery was ligated for 35 min. After that, the mice were placed in a nursing box at 37°C for 15 minutes for recovery and then returned to their home cages.

Three days after modeling, the mice were killed by quick decapitation and transferred to an ultraclean worktable for dissection. Then, 75% alcohol was used to clean the abdomens, and an incision was made in the abdominal cavity to completely separate the intestines. One to two mouse feces were collected from the end of the colon and stored in a sterilized EP tube. The feces were rapidly frozen and stored in a freezer at −80°C.

### 2.4 Mouse droppings sequencing experiment

Mouse fecal samples were removed from the −80°C freezer and then immediately sent to Tianhao Biotech Company (Shanghai) on dry ice for sequencing. The sequencing procedure was as follows: an excrement gene DNA extraction kit was used to extract mouse fecal DNA, and after the quality of the DNA verified by 1% agarose gel electrophoresis, polymerase chain reaction (PCR) was performed. Three identical PCR experiments were performed for each sample. The v3-v4 region of the 16S rRNA gene was selected to be amplified to synthesize a fusion primer with dislocation bases. The number of amplification cycles of the samples was guaranteed to be consistent, and the majority of samples was amplified to show products at the appropriate concentration. The three PCR products of the same sample were mixed and detected by 2% agarose gel electrophoresis, and then the PCR products were recovered using an AxyPrep DNA gel extraction kit. The PCR products were quantified by an enzyme marker and mixed in proportion. After a MiSeq PE library was constructed, an Illumina MiSeq PE 300 platform was used for high-throughput sequencing.

### 2.5 Biological information analysis

TrimGalore software was used to remove the bases with a terminal mass of quality less than Q20 and to remove the adapter sequences that might be included. Then, short sequences with a length of less than 100 bp were removed. Using FLASH2 software to splice the paired sequences obtained by double-terminal sequencing to obtain the merged sequence, the low-quality sequences were further removed after merging. Then, mothur software was used to find and remove primers in the sequence, and Usearch software was used to remove sequences with which the total base error rate was greater than Q20 or the sequence length was less than 100 bp to obtain clean reads with high quality and reliability, which were then used for subsequent bioinformatics analysis. Qiime (v1.8.0) software was used for operational taxonomic unit (OTU) clustering; species annotations with greater than 97% similarity were assorted to an OTU. The representative OUT sequences were compared with the corresponding microbial database to determine the species classification of each sample, and OTUs were annotated at all levels.

### 2.6 Community composition histogram and diversity analysis

Alpha diversity analysis, community composition analysis and beta diversity analysis were carried out based on the clustering results. We used R software (version 3.6.2) to perform Alpha diversity analysis, community composition analysis and Beta diversity analysis. We used the R packages microbiome (version 1.8.0) and microbiomeseq (version 0.1) to complete the analysis.

### 2.7 SCFA preparation and extraction

Fecal samples were weighed, 20 mg of each sample was placed in a 2-ml EP tube, and 1 mL of phosphoric acid (0.5% v/v) was added to each EP tube. The tubes were then vortexed for 10 min, and ultrasonicated for 5 min. Then, 0.1 mL of supernatant was added to each 1.5 mL centrifugal tube, 0.5 mL of MTBE (containing the internal standard) was added, and the tubes were then vortexed for 3 minutes and ultrasonicated for 5 minutes. After that, the tubes were centrifuged for 10 minutes at 12,000 r/min and 4°C. After centrifugation, 0.2 mL of supernatant was absorbed into the sampling bottle for GC-MS/MS analysis.

## Results

### 3.1 Data quality control and statistics

The data quality is shown in Table 2.

**Table 2.** Mean data quality of the samples

|  | Raw_reads | Q20<br>(%) | Q30<br>(%) | Merge | Merge<br>(%) | Primer | Clean_reads | Clean_reads<br>(%) | Q20<br>(%) | Q30<br>(%) |
| --- | --- | --- | --- | --- | --- | --- | --- | --- | --- | --- |
| Samples | 233087.8 | 98.05 | 94.19 | 230179 | 98.75 | 229118 | 214464.2 | 92.02 | 99.21 | 96.63 |

The Q20% of raw reads was 98.05%, and the Q30% of raw reads was 94.19%. After removing the adapter sequences and primers from the sequences and filtering low-quality sequences, we obtained clean reads. The Q20% of the raw reads was 99.21%, and the Q30% of raw reads was 96.63%, which met the quality requirements.

### 3.2 Alpha diversity

Rarefaction curve analysis (Fig. 1) showed that as the number of extracted sequences increased, it reached a stable level, indicating that the sequencing depth had covered rare new phylotypes and most of the diversity. In addition, the numbers of OTUs were different between groups. The sham germ-free groups (SGFCG, SGFEG, SGFNG) had fewer kinds of OTUs than the other groups, which demonstrated that it is possible to construct sham germ-free models treated with antibiotics.

**Fig. 1.**
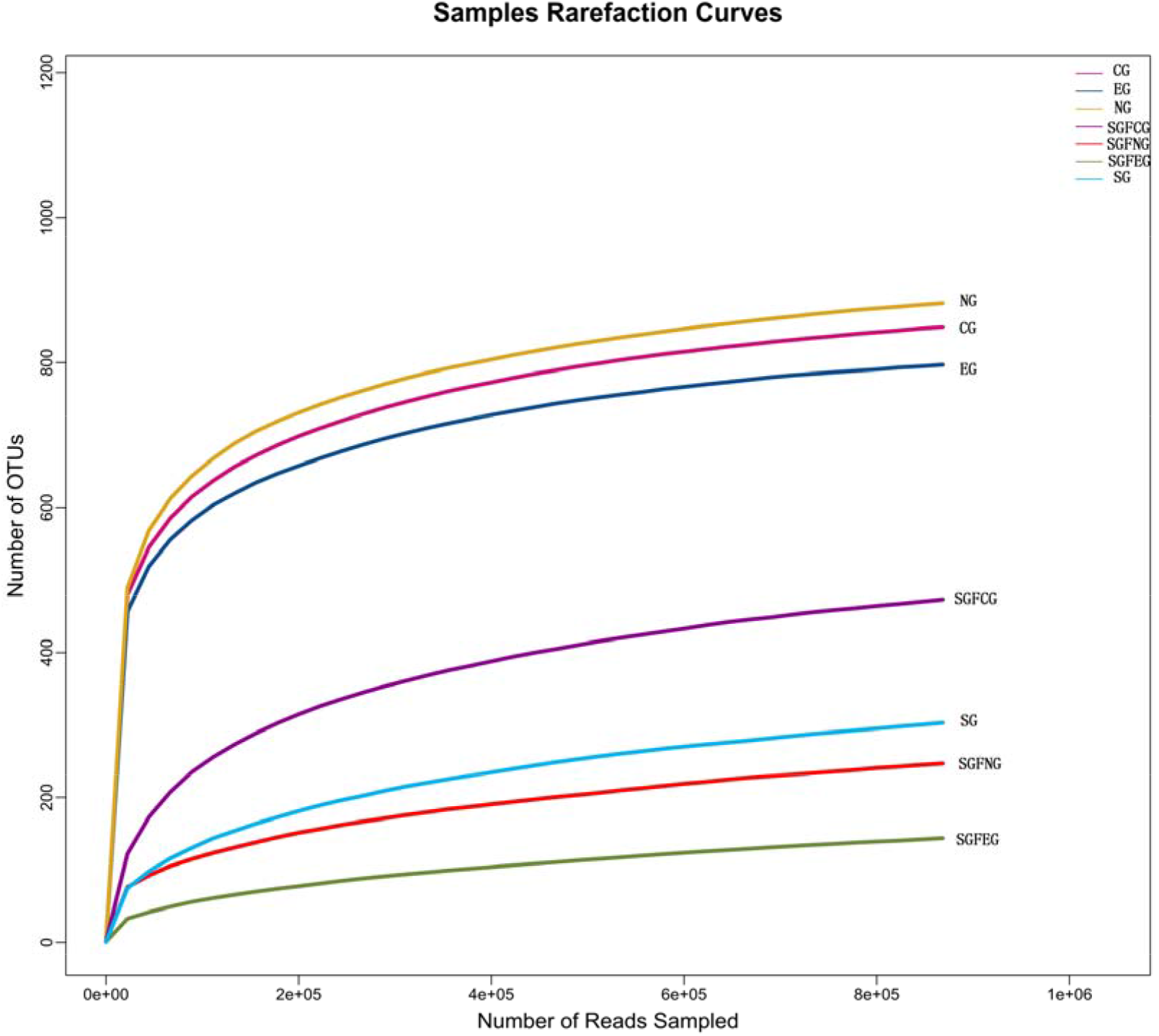
Rarefaction curves for all groups.

Gut microbiota diversity and richness were evaluated by the species richness index, Simpson index and Shannon index. The results between the CG, SGFCG and EG, SGFEG groups are shown in Fig. 2, and the results of all of groups are shown in Fig. 3.

**Fig. 2.**
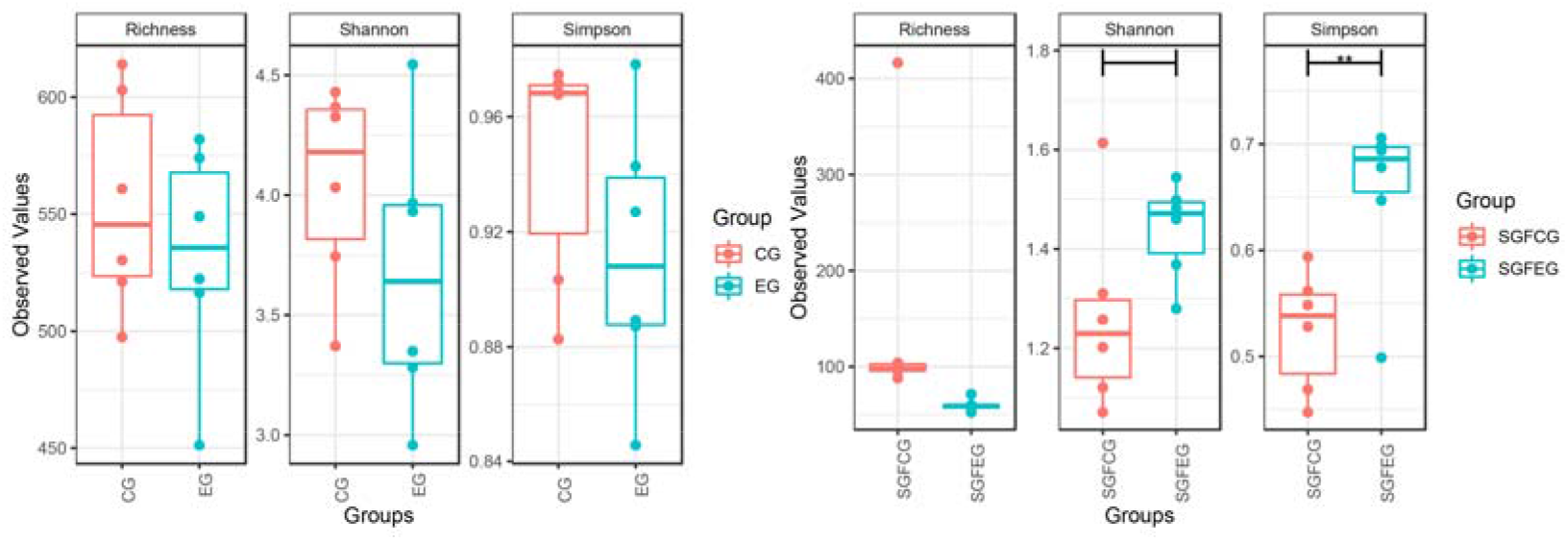
Richness, Simpson and Shannon indexes between the XNJ and control groups (∗∗, P<0.01 between the two groups).

**Fig. 3.**
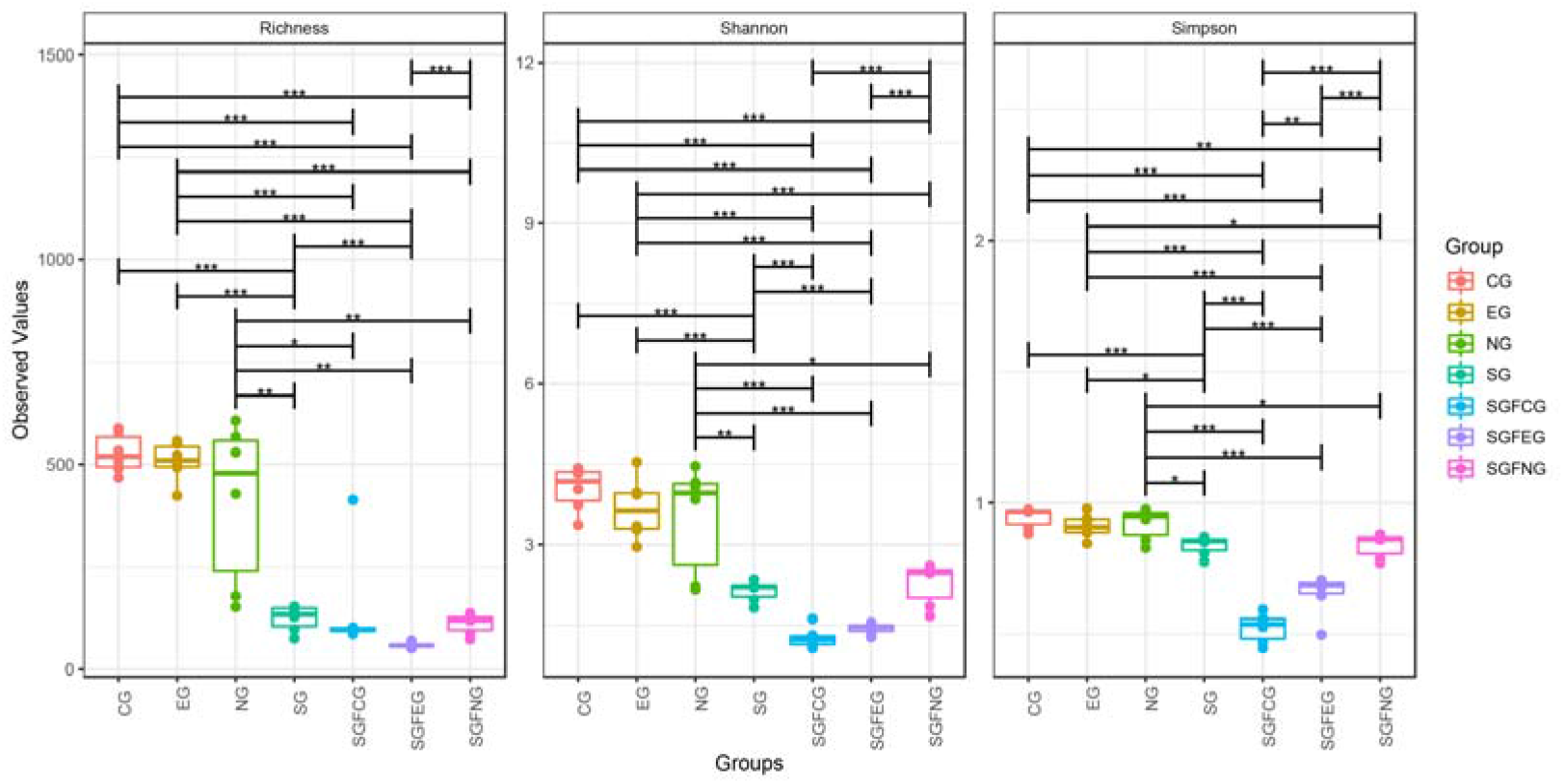
Richness, Simpson and Shannon indexes in all groups (*, P<0.05, **, P<0.01, ***, P<0.001 between the two groups).

The differences in diversity and richness between the CG and EG were not significant. The Simpson index of the SGFEG was significantly greater than that of the SGFCG, but the richness of the SGFCG was greater than that of the SGFEG. When all groups were compared, there were significant differences between the sham germ-free group and the other groups, with lower alpha diversity in the sham germ-free group. Moreover, there were no significant difference in alpha diversity between the model and normal group.

### 3.3 Beta analysis

We used several different algorithms to try to separate groups. PCoA, DPCoA, DCA, CCA, NMDS, MDS, and RDA was performed to compare the overall microbiota structures in each group (Fig. 4).

**Fig. 4.**
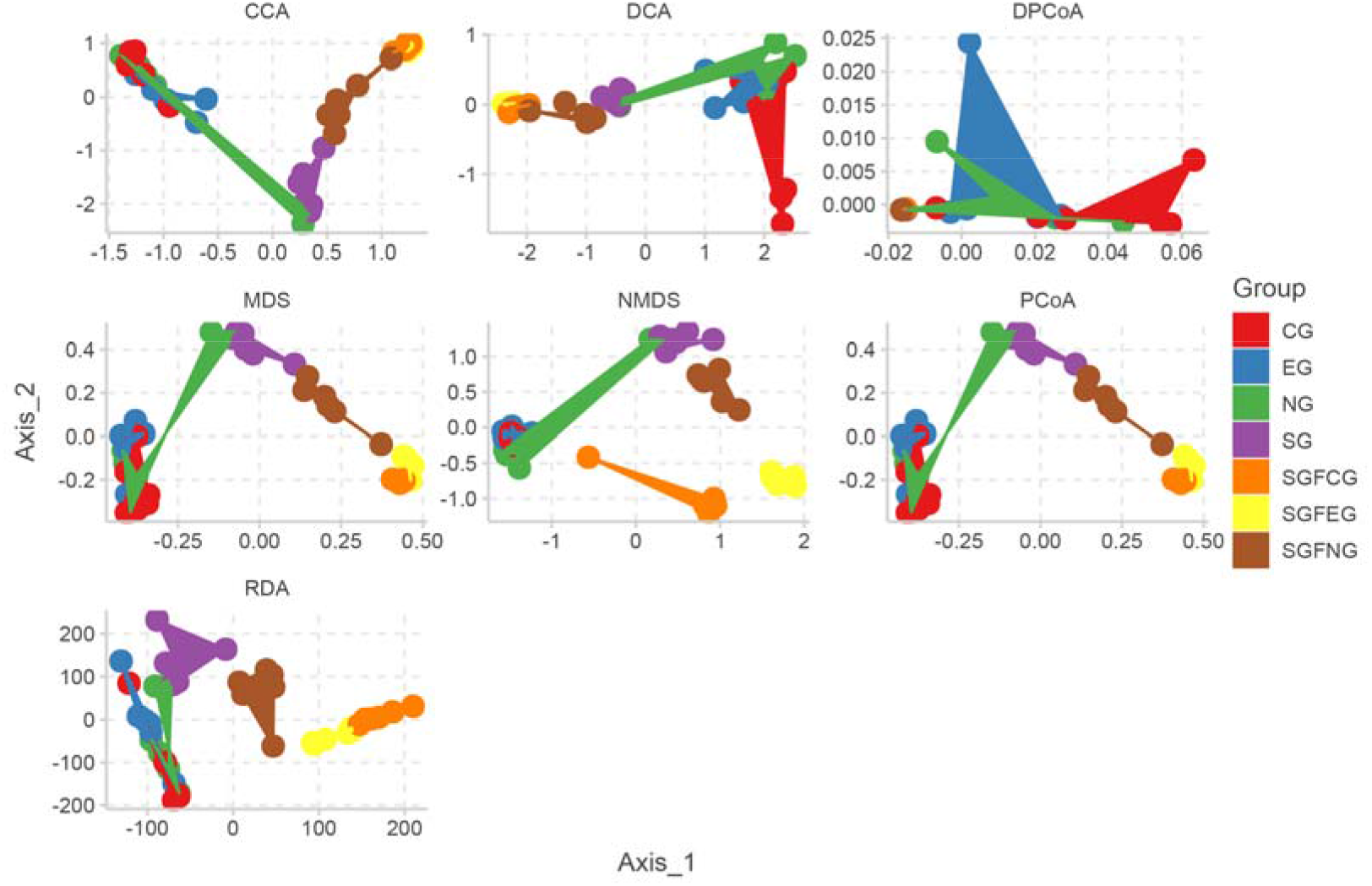
PCoA, DPCoA, DCA, CCA, NMDS, MDS, and RDA for all groups.

CG and EG clusters could be divided into two groups by DPCoA. For most algorithms, the SGFCG and SGFEG could be divided. These results suggested that XNJ injection changed the overall gut microbiota composition between the CG, SGFCG and EG, SGFEG groups and that these group could be divided by some algorithms.

We also performed LEfSe analysis to identify the specific bacterial genera that were characteristic among the 7 groups. The results are shown in Fig. 5.

**Fig. 5.**
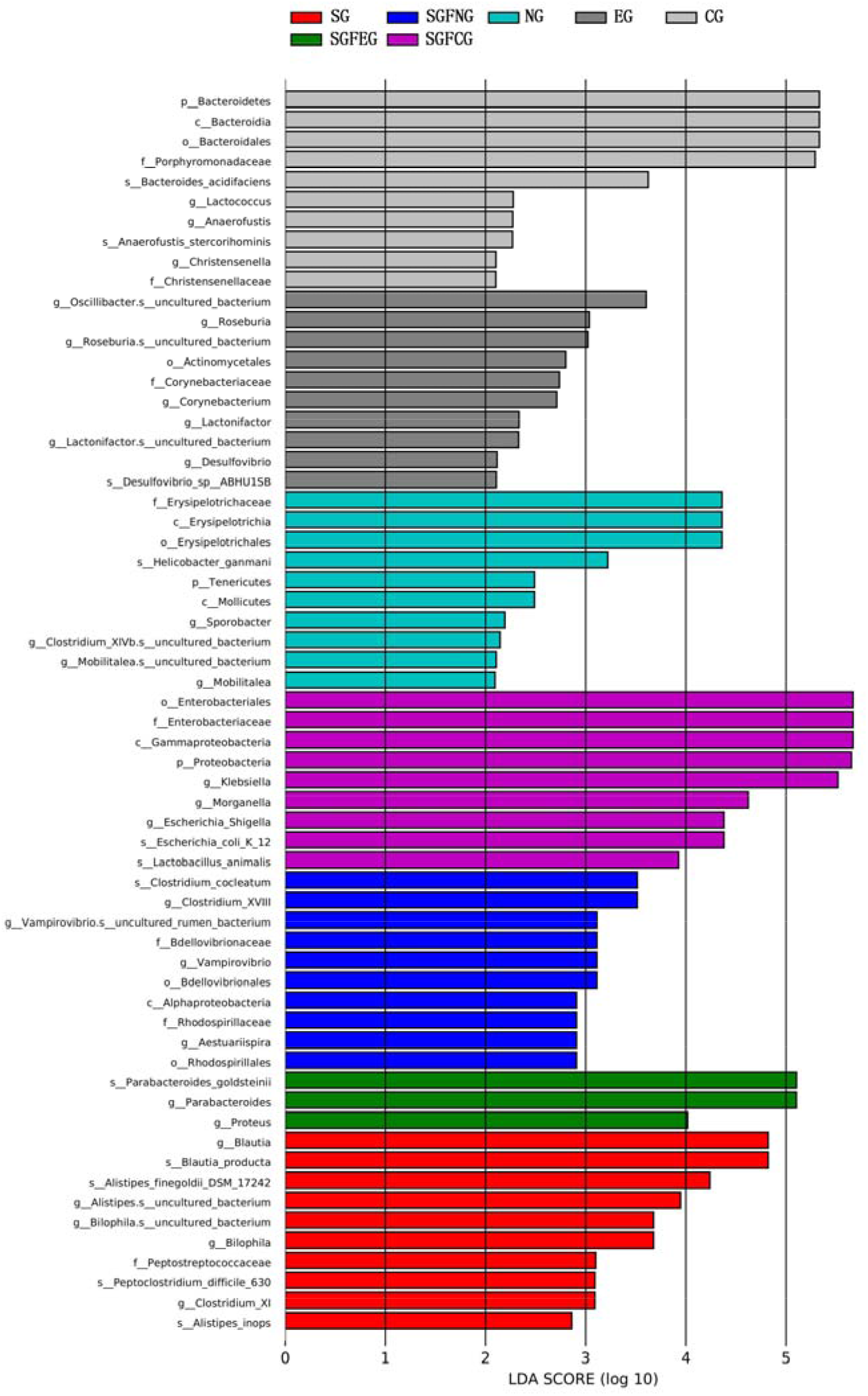
LEfSe analysis at the genus level.

### 3.4 Community composition analysis

We compared community composition between the SGFCG and SGFEG, EG and CG at the phylum, family and genus levels to identify the primary types of gut microbiota that were differently expressed between the CG, SGFCG and EG, SGFEG groups.

At the family level, as indicated in Fig. 6A, while the EG and CG group mice and the SGFCG and SGFEG group mice presented very similar gut microbiota compositions, XNJ significantly affected the relative abundances of certain families. The differences in the gut microbiota at the family level are shown in Fig. 6B and Fig. 6C. Specifically, there were significantly differences in *Bacteroidaceae*, *Rikenellaceae*, *Ruminococcaceae*, *Erysipelotrichaceae*, *Moraxellaceae* and *Sutterellaceae* between the SGFCG and SGFEG, with *Sutterellaceae* being a dominant microbiota family in the SGFCG and SGFEG. *Enterobacteriaceae*, *Peptostreptococcaceae*, *Flavobacteriaceae* and *Deferribacteraceae* were significantly different between the CG and EG, while none of these families were dominant in CG and EG. *Enterobacteriaceae*, *Lachnospiraceae*, *Porphyromonadaceae*, *Verrucomicrobiaceae*, *Bacteroidaceae* and *Ruminococcaceae* were dominant families in all of the samples. We also found that there were robust differences between germ-free mice and normal mice. In addition to the above results, we found that Xingnaojing regulated the abundance of *Sutterellaceae* in the SGFCG and SGFEG.

**Fig. 6.**
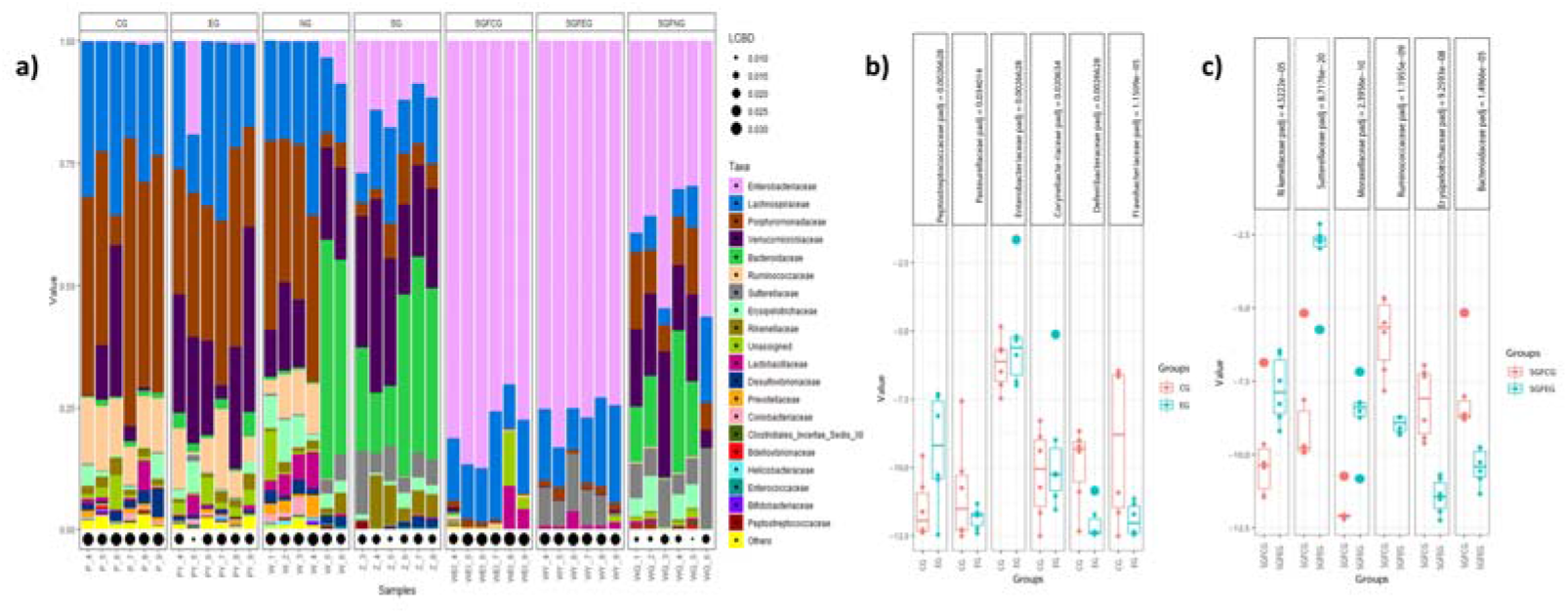
Community composition at the family level and significantly different bacteria with adjusted p-values. a) Community compositions at the family level in all groups. b) Significantly different bacteria between the CG and EG. c) Significantly different bacteria between the SGFCG and SGFEG. (we set the p-adjusted threshold between the SGFCG and SGFEG to 0.001 and the p-adjusted threshold between the EG and CG to 0.05).

At the genus level, the different kinds of gut microbiota were as follows (Fig. 7B and Fig. 7C). The components of the gut microbiota at the genus level are shown in Fig. 7A. There were robust differences between *Morganella Parasutterella* between the SGFCG and SGFEG; *Morganella* levels decreased after Xingnaojing injection, while *Parasutterella* levels increased. Akkermansia was the main component of the gut microbiota in the EG, and there were some differences between the CG and EG (p-adjust=0.45). *Morganella* was downregulated in both groups.

**Fig. 7.**
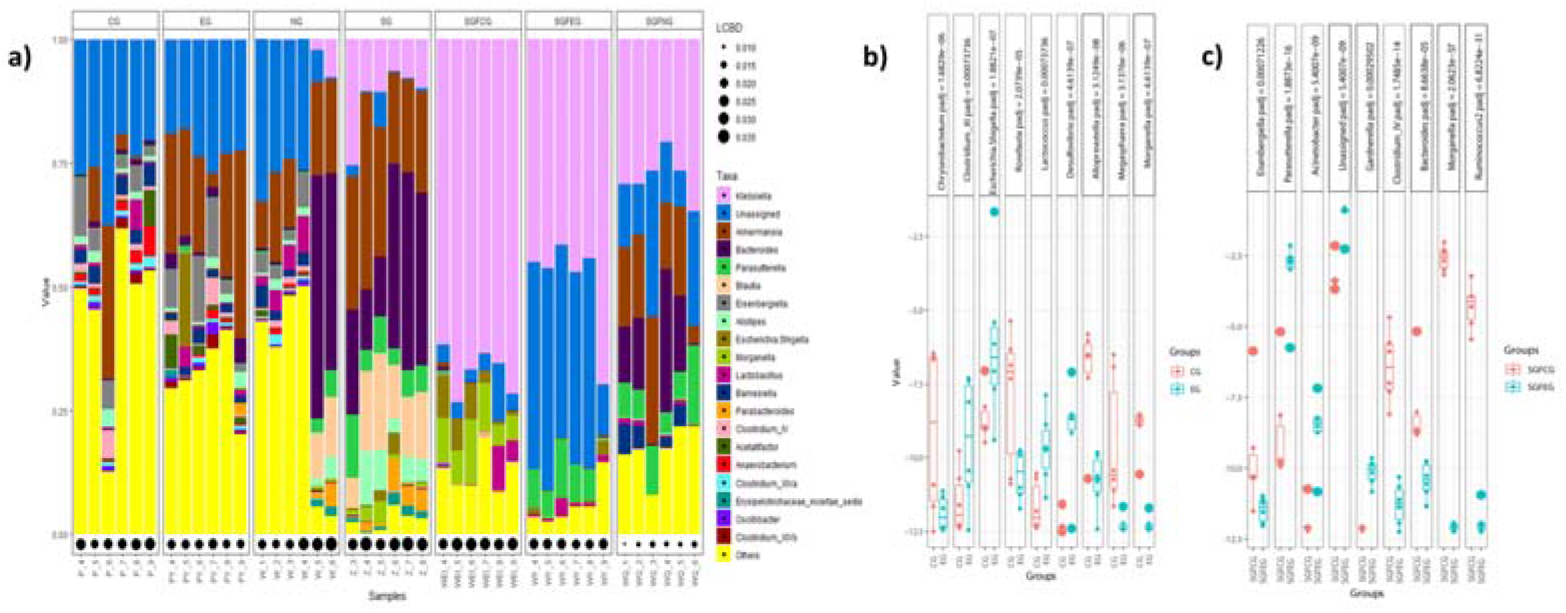
Community composition at the genus level and significantly different bacteria with p-adjusted values. a) Community compositions at the genus level of all groups. b) Significantly different bacteria between the CG and EG. c) Significantly different bacteria between the SGFCG and SGFEG. (we set p-adjusted threshold between the SGFCG and SGFEG to 0.001 and the p-adjusted threshold between the EG and CG to 0.001).

At the phylum level, two kinds of bacteria were different between the CG and EG, while one kind of bacteria was different between the SGFCG and SGFEG (p-adjusted <0.15). Firmicutes, *Bacteroidetes* and *Verrucomicrobia* were most common components of the gut microbiota in CG and EG. In the SGFCG and SGFEG groups, *Proteobacteria*, *Firmicutes* and *Bacteroidetes* were most common. *Deferribacteres* levels were decreased in both groups (Fig. 8).

**Fig. 8.**
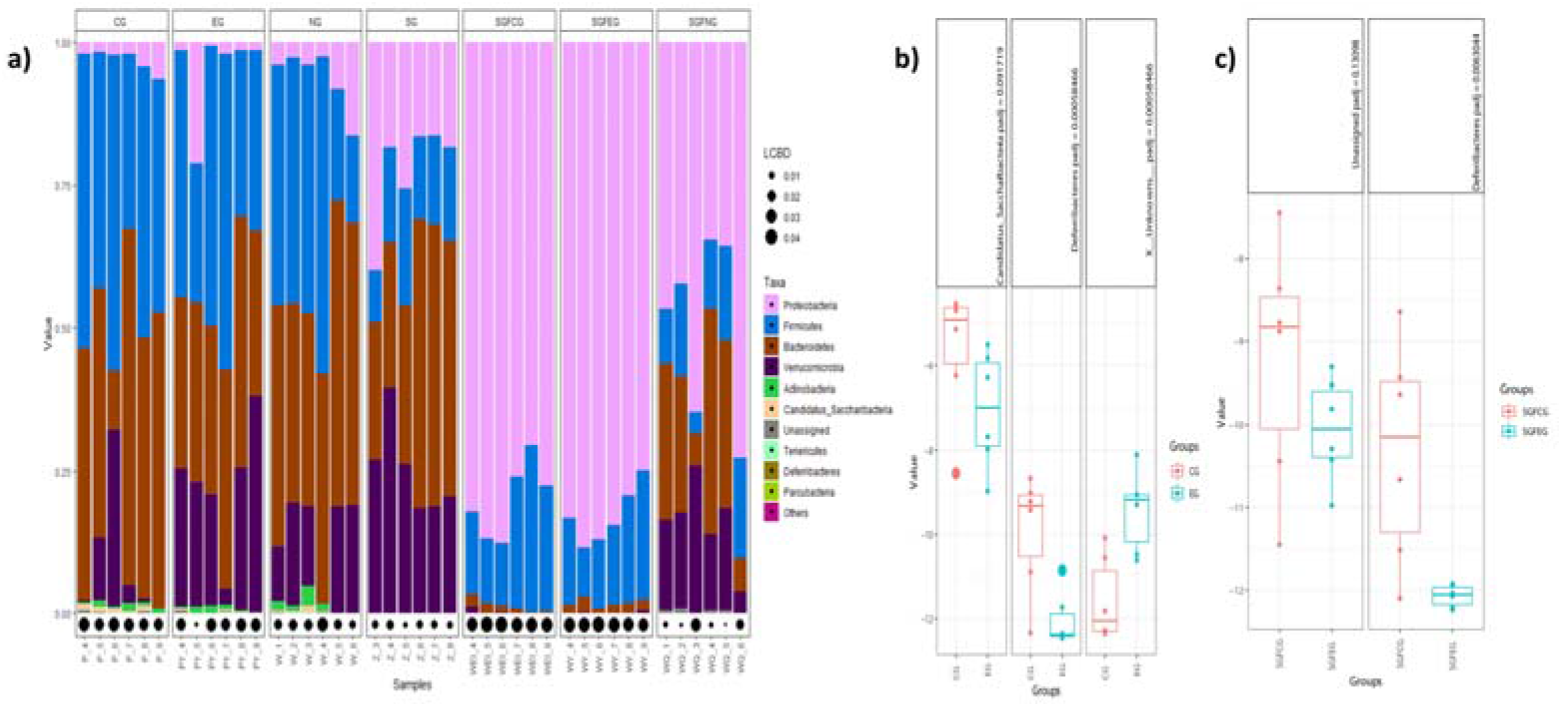
Community composition at the phylum level and significantly different bacteria with p-adjusted values. a) Community compositions at the phylum level in all groups. b) Significantly different bacteria between the CG and EG. c) Significantly different bacteria between the SGFCG and SGFEG. (We set the p-adjusted threshold between the SGFCG and SGFEG to 0.15 and the p-adjusted threshold between the EG and CG to 0.15),

### 3.5 Functional prediction

We used PICRUSt to predict the functions of gut bacteria, and differentially enriched KEGG pathways between the XNJ (EG and SGFEG) and control groups (CG and SGFCG) were identified (Fig. 9). We found that peptidoglycan biosynthesis was most different between CG and EG. Lipoic acid metabolism, antigen processing and presentation, progesterone-mediated oocyte maturation, and transporters were the most different between the SGFCG and SGFEG.

**Fig. 9.**
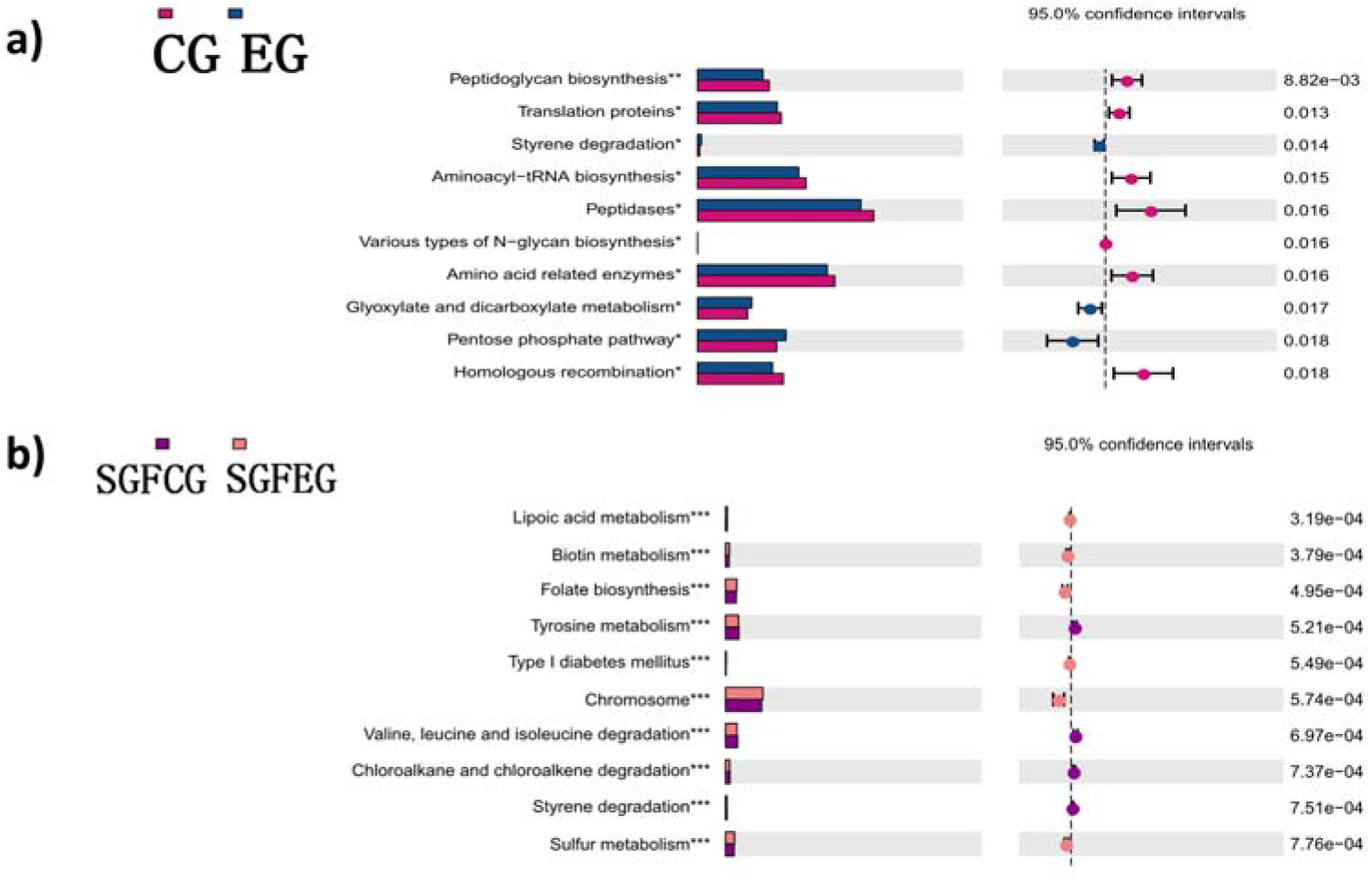
Differential enrichment of KEGG pathways. a) Differential enrichment of KEGG pathways between the CG and EG. b) Differential enrichment of KEGG pathways between the SGFCG and SGFEG (*, P<0.05, **, P<0.001, ***, P<0.0001 between the two groups).

### 3.6 Analysis of SCFAs in the feces

We evaluated the levels of SCFAs in feces for all groups (Fig. 10). We found that the SCFA levels in the sham germ-free groups were lower than those in the other groups. Moreover, XNJ increased the levels of PA, VA, IBA, and IVA in the SGFEG group (t test, p-value<0.05). SCFA levels were not significantly different between the EG and CG.

**Fig. 10.**
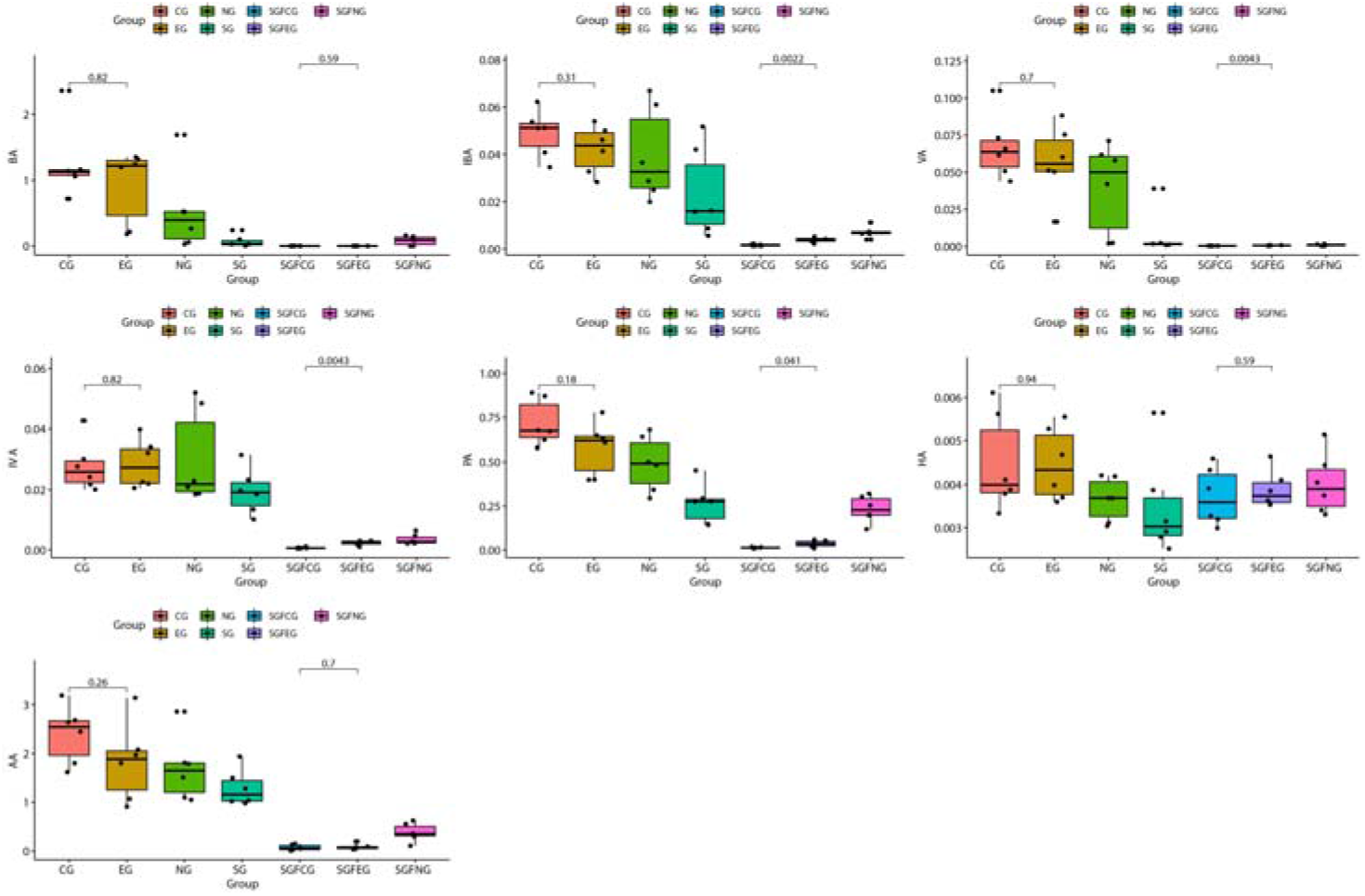
Plots of the SCFA levels in the feces in all groups (VA: valeric acid/valerate, IVA: isovaleric acid/isovalerate, IBA: isobutyric acid/isobutyrate, HA: caproic acid/caproate, AA: acetic acid/acetate, PA: propionic acid/propionate, BA: butyric acid/butyrate); p-values between the CG and EG and between the SGFCG and SGFEG are indicated in the plots.

### 3.7 Correlation between SCFA levels and the gut microbiota

We performed single-factor regression analysis for all kinds of microbiota at the genus level and screened for p-values<0.00001. The results showed that at the genus level, 8 kinds of bacteria (*Saccharibacteria_genera_incertae_sedis*, *Klebsiella*, *Intestinimonas*, *Oscillibacter*, *Aestuariispira*, *Anaerotruncus*, *Eisenbergiella*, and *Clostridium_XlVa*) had a strong possibility of having a linear relationships with the SCFA concentration (Fig. 11a). We performed Pearson correlation analysis on the 8 kinds of bacteria and SCFAs, and the correlation coefficients are shown in Fig. 11b. Most absolute values of correlation coefficients between SCFA levels and the 8 kinds of bacteria were greater than 0.5.

**Fig. 11a:**
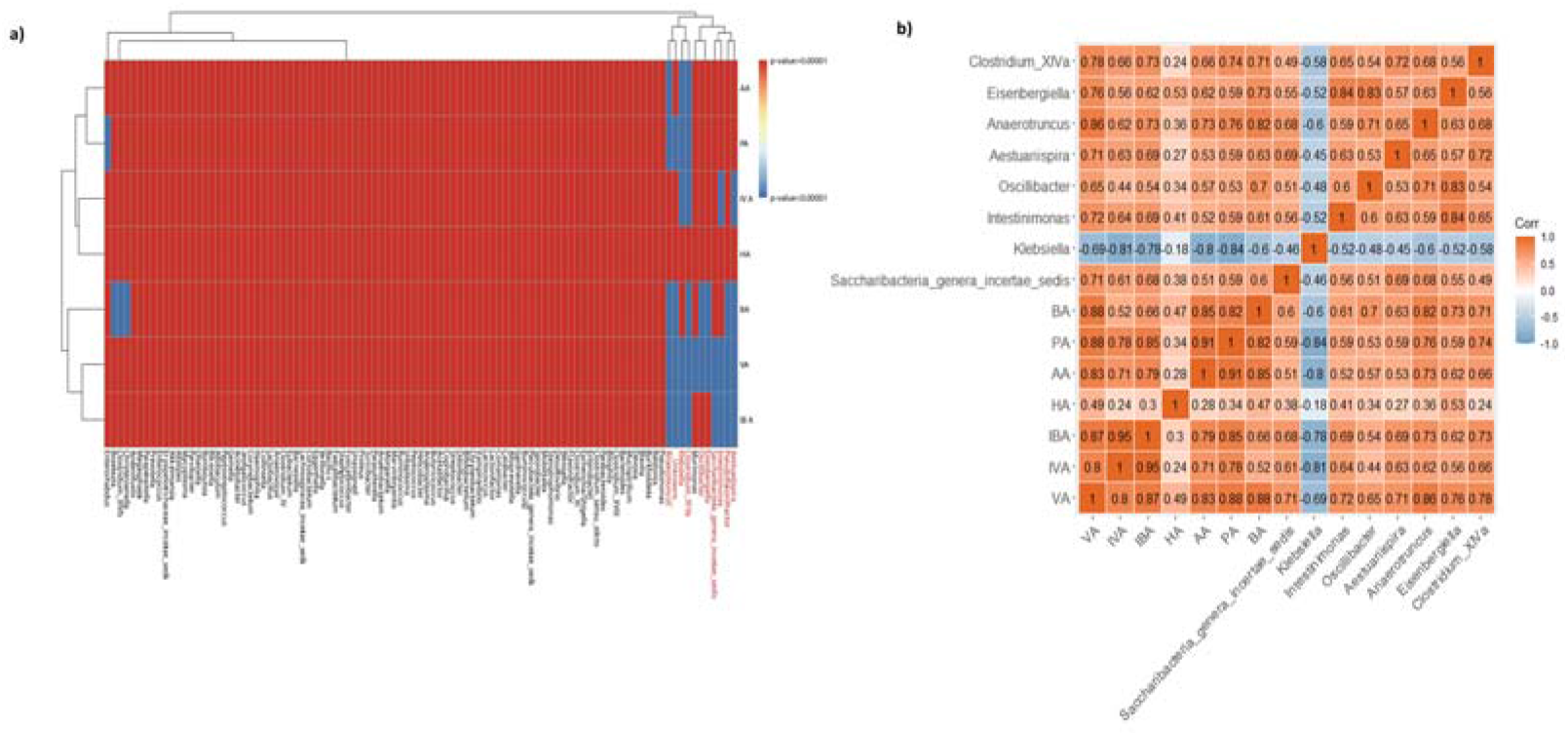
Single-factor regression analysis for all kinds of bacteria at the genus level with a p-value<0.00001 (red: P>0.00001; blue: P<0.00001). Fig. 11b: Pearson correlation analysis of the 8 selected bacteria (red: positive correlation; blue: negative correlation; shade of color: absolute value of correlation coefficients).

We plotted the correlations between SCFA levels and the gut microbiota at the phylum and genus levels for all groups (Fig. 10 and Fig. 11). We determined that many kinds of bacteria (*Verrucomicrobia*, *Tenericutes*, *Actinobacteria*, and *Candidatus_Saccharibacteria* at the phylum level and many more kinds of bacteria at the genus level) exhibited strong positive Pearson correlations with butyrate levels in the SGFCG group, which were significantly different from those in the other groups. XNJ changed this trend, resulting in no obvious correlation between gut bacteria and butyrate levels in the SGFEG group. Further analysis indicated that the bacteria that were strongly correlated with butyrate levels at the phylum level were in low abundance in the SGFCG and SGFEG (Fig. 13).

## Discussion

Ischemic stroke (IS) is the most common type of stroke and is caused by thrombotic or embolic occlusion and stenosis of a cerebral artery due to sudden loss of blood circulation to an area of the brain, resulting in a corresponding loss of neurologic function. It has a high incidence, high disability rate and high mortality rate^[23, 24]^. We generated a middle cerebral artery occlusion (MCAO) mouse model as a model of IS to explore the mechanism underlying the effect of XNJ injection on IS from the perspective of the intestinal microbiota.

The quality of sequencing data was very good, as the Q30 of the raw data was greater than 90%, and that of the clean data was greater than 95%.

Community composition and alpha diversity analyses showed that XNJ injection modulated the overall structure of the gut microbiota in C57 mice and resulted in large differences between the sham germ-free groups and the other groups.

Regarding alpha diversity, we found that the difference in diversity was not significant in the CG and EG but that the diversity of the EG was lower than that of the CG overall. Previous studies have indicated that IS patients and IS models exhibit no significant differences in alpha diversity compared with the control group[20, 25, 26]. A previous study of the effect of the combination of the Chinese medicines Puerariae Lobatae Radix and Chuanxiong Rhizoma on the mechanism of gut microbiota in cerebral ischemic stroke found that the injection group had a significantly decreased diversity compared with that of the control group^[27, 28]^ and that the trends in Alpha diversity were in accordance with those observed between the CG and EG. However, in the SGFCG and SGFEG, the diversity of the XNJ group was greater than that of the control group, which contradicted the results in obtained for the CG and EG, indicating that XNJ can decrease the high diversity in the CG and increase the low diversity in the SGFCG. We hypothesized that XNJ can bidirectionally regulate the alpha diversity of the gut microbiota, which should be further researched.

We found that XNJ increased the level of *Sutterellaceae* in the SGFEG compared with the SGFCG. *Sutterellaceae* is mainly found in the intestinal tract of humans and some animals as a member of the indigenous intestinal microbiota and can be isolated from both the intestinal tract and from infections of gastrointestinal origin^[29]^. At the genus level, *Morganella* was significantly increased in the SGFCG group compared to the SGFEG group and decreased after XNJ injection gavage, while *Parasutterella* was increased. In previous studies, we also found that *Morganella* was significantly increased in IS patients^[30]^. *Morganella* is a genus of Gram-negative bacteria that has a commensal relationship within the intestinal tracts of humans, mammals, and reptiles as a member of the normal flora and can be a pathogen in nosocomial infections^[31]^. *Parasutterella* can fill an ecological niche in the gastrointestinal tract and contribute to metabolic functionalities^[32]^. At the phylum level, *Deferribacteres* was decreased in both the EG and SGFEG. In Runzhi Chen’s study on Puerariae Lobatae Radix and Chuanxiong Rhizoma^[27]^, the Chinese medicines could also regulate the gut microbiota by reducing *Deferribacteres*, a phylum of gram-negative bacteria that makes energy through anaerobic respiration^[33]^. LEfse analysis revealed that SGFCG mice were enriched with *Morganella*, which is in agreement with previous research^[30]^. XNJ also decreased this trend in the SGFEG group.

KEGG analysis showed that peptidoglycan biosynthesis was most different between the CG and EG; for map00550, we found that peptidoglycan is a macromolecule made of long amino-sugar strands cross-linked by short peptides. It forms the cell wall in bacteria surrounding the cytoplasmic membrane[34]. In the SGFCG and SGFEG, we found that lipoic acid metabolism and boitin metabolism were most enriched, which function mainly in the synthesis and metabolism of organic matter.

XNJ increased the levels of PA (propionate), VA (valerate), IBA (isobutyrate), and IVA (isovalerate) in the feces of the SGFEG group. SCFAs are generated by bacterial fermentation of polysaccharides^[35]^, and the composition of an individual’s microbiota and the presence of keystone species also influence fiber fermentation^[36]^. Thus, we concluded the XNJ increased the community diversity in the SGFEG to induce this phenomenon. Shannon Rose^[37]^ suggested that bacterial metabolites (SCFAs) may affect mitochondrial function to elicit mitochondrial dysfunction-induced the cecal gut dysmotility and that mitochondrial dysfunction can cause gut dysmotility. For example, butyrate is converted into acetyl-CoA, which is then utilized in the citric cycle for NADH production, and NADH is an important substance for mitochondrial function[38]. Therefore, we believe that XNJ can improve IS prognosis, partly by regulating SCFA levels in the intestine.

We screened 8 kinds of bacteria that had a high possibility of having a linear relationship with the concentration of SCFAs. Previous research has revealed that SCFAs are produced by several bacteria from different pathways^[39–41]^. However, only *Clostridium* are involved in producing butyrate. As much of the information on the diversity of SCFA-producing bacteria has depended on culture methods[42, 43] and the culture methods were difficult to found new kinds of SCFA-producing bacteria, we hypothesized that some of the screened bacteria *(Saccharibacteria_genera_incertae_sedis*, *Klebsiella*, *Intestinimonas*, *Oscillibacter*, *Aestuariispira*, *Anaerotruncus*, *Eisenbergiella*, and *Clostridium_XlVa*) generate or inhibit SCFAs. We found that butyrate had a greater positive correlation with gut bacteria than other SCFAs in the SGFCG and that XNJ downregulated this trend, resulting in no obvious correlation between gut bacteria and butyrate levels (Fig. 12A and Fig. 12B). Some studies have also focused on the changes in SCFA levels in patient feces^[38, 44–49]^. *Eubacterium*, *Clostridium*, *Faecalibacterium prausnitzii*, and *Roseburia* are thought to be butyrate-producing bacteria^[39, 48, 49]^; however, as *Eubacterium*, *Faecalibacterium prausnitzii* and *Roseburia* were low in abundance at the genus level, they are not shown in Fig. 12B. However, low abundances of *Clostridium XlVa* were found in the SGFEG and SGFCG (Fig. 12D); thus, we hypothesize that the contribution of *Clostridium XlVa* was small.

**Fig. 12a:**
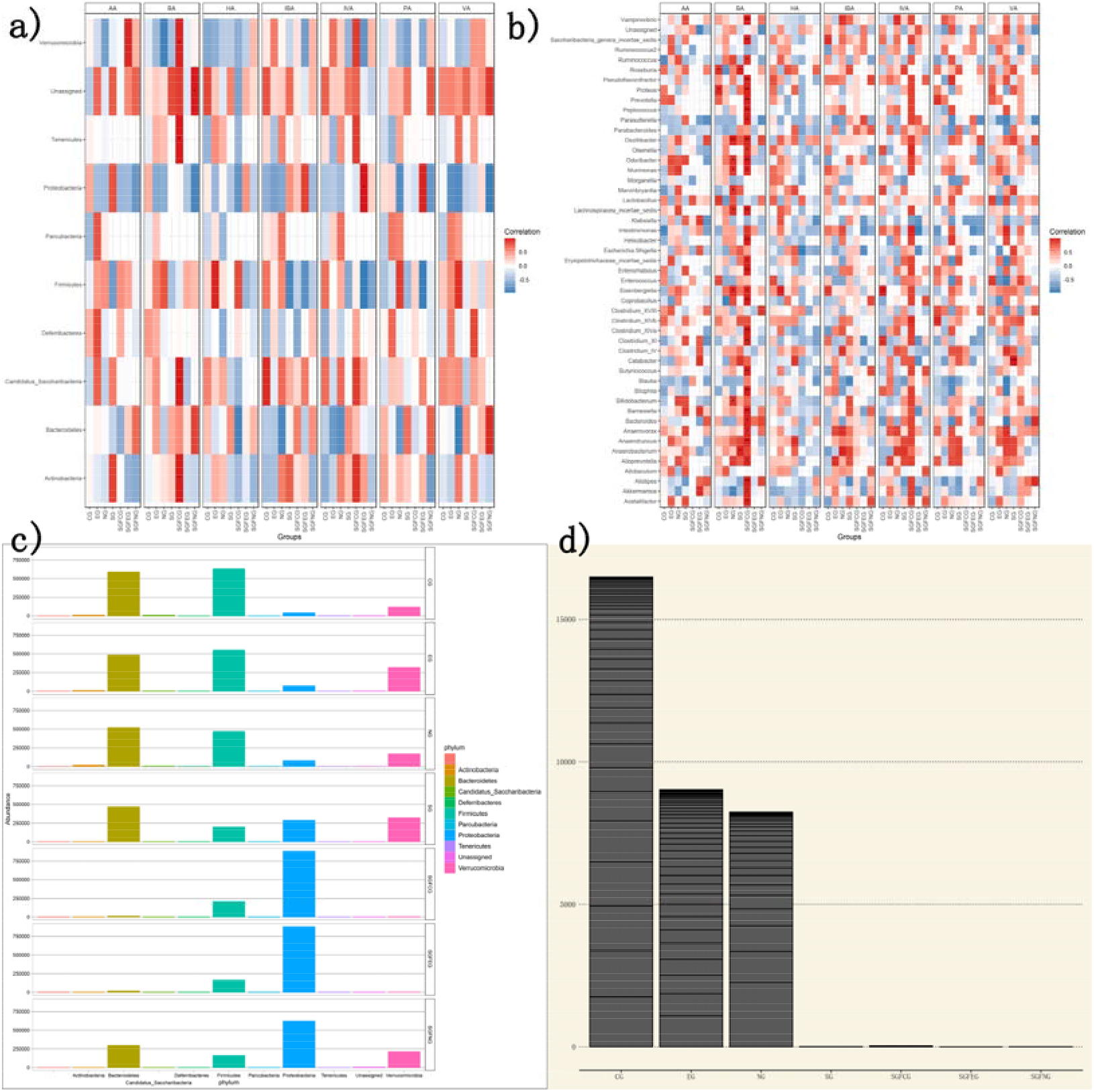
Correlations between SCFA levels and the gut microbiota at the phylum level (*P<0.05, **P<0.01). Fig. 12b: Correlations between SCFA levels and the gut microbiota at the genus level (*P<0.05, **P<0.01). Fig. 12C: Bar graph of the abundance of the gut microbiota at the phylum level. Fig. 12D: Bar graph of the abundance of *Clostridium XlVa* at the genus level.

Other research has indicated that a host genetics-driven increase in the gut production of butyrate is associated with improvements in metabolic traits[47]. Therefore, the causes of the positive correlation between BA levels but not other SCFAs and gut bacteria in the SGFCG should be further researched. We hypothesize that as the diversity of gut microbiota decreases in the SGFCG, the butyrate level decreases, but some mechanisms may occur after XNJ injection, disrupting this correlation. The abundances of correlated bacteria were very low in the SGFCG and SGFEG at the phylum (Fig. 12C) and genus levels; thus, the butyrate level did not change much overall (Fig. 10). More attention should also be paid to the subtle mechanism by which XNJ injection changes the correlation between specific gut bacteria and butyrate levels.

Based on the overall results of the experiment, we found that the SGFCG and SGFEG exhibited much greater differences in alpha diversity and SCFA levels than the CG and EG. We hypothesize that oral administration of antibiotics decreases the richness of gut bacteria, which may control the test variables. On the other hand, the CG and EG exhibited more complex results, which may negate their differences compared to the SGFCG and SGFEG. However, the CG and EG might simulate human ischemic stroke conditions better than the SGFCG and SGFEG. Germ-free mouse models are generally considered to be the gold standard for studies of the microbiota^[50]^. The use of antibiotic treatments to construct a sham germ-free group is an alternate method. Mice on antibiotics are not completely cleared of bacteria but exhibit significant reductions in bacterial load^[50]^, which could control variables and result in more obvious results.

Because of the short nature of the experiment, the functions of selected different bacteria should be further observed, possibly by planting specific bacteria to explore their functions. Second, the KEGG pathways of the functions were only calculated by OTUs; verification should also be performed at the biochemical and physiological levels. Third, regarding SCFAs, the correlation between the gut bacteria and butyrate levels after XNJ injection should be further explored at the biochemical and physiological levels.

## 5. Conclusions

In conclusion, ischemic stroke causes dysbiosis of some specific bacteria in the gut microbiota. Xingnaojing can ameliorate this condition by increasing the levels of *Sutterellaceae* and decreasing the level of *Deferribacteres* and *Morganella*. These results are in accordance with other research on the use of Chinese medicines for that affect the gut microbiota for IS treatment. The results of the SCFA analysis and enrichment analysis revealed that XNJ regulated the gut microbiota and short-chain fatty acids, mainly through energy metabolism-relevant pathways. Moreover, the specific mechanisms underlying the XNJ injection-induced change in the gut microbiota and short-chain fatty acids in IS should be further researched.

## List of abbreviations

XNJ: Xingnaojing injection
MCAO: middle cerebral artery occlusion
SCFA: short-chain fatty acid
IS: ischemic stroke
AIS: acute ischemic stroke
GBA: gut-brain axis
TMAO: trimethylamine N-oxide
OUT: operational taxonomic unit
VA: valeric acid/valerate
IVA: isovaleric acid/isovalerate
IBA: isobutyric acid/isobutyrate
HA: caproic acid/caproate
AA: acetic acid/acetate
PA: propionic acid/propionate
BA: butyric acid/butyrate

## Declarations

### Ethics approval and consent to participate

This research was approved by Dongzhimen Hospital Ethic Committee, Beijing University of Chinese Medicine.

### Consent for publication

Written informed consent for publication was obtained from all participants.

### Availability of data and materials

The datasets used or analysed during the current study are available from the corresponding author on reasonable request.

### Authors contributions

J.L., G.L., Q.G., Z.W. and Z.C. performed the biological experiment. J.L. and G.L. performed data analysis. D.M., Y.G. and Z.H. conceived and supervised the study. J.L. and D.M. drafted the manuscript. All authors read and approved the final manuscript.

## Acknowledgements

Thanks to AJE for native English polish.

